# The global phylogeography of rapidly expanding multidrug resistant Ural lineage 4.2 *Mycobacterium tuberculosis*

**DOI:** 10.1101/2025.08.13.670097

**Authors:** Melanie H. Chitwood, Isabel Rancu, Yexuan Song, Barney Potter, Yi Ting Chew, Nelly Ciobanu, Valeriu Crudu, Caroline Colijn, Ted Cohen, Benjamin Sobkowiak

## Abstract

Multidrug resistant tuberculosis (MDR-TB) is a leading global health challenge. Recent studies identified a reproductively fit Ural lineage 4.2 MDR *M. tuberculosis* (*Mtb*) variant circulating within the Republic of Moldova. We searched a large publicly available dataset of ∼200,000 *Mtb* whole genome sequences and identified a clade of 1604 drug-resistant *Mtb* sequences harboring conserved resistance-conferring mutations that are closely related to the MDR *Mtb* variant circulating in Moldova. We identified the Russian Federation as the most likely country of origin for this clade, and we inferred several independent migration events from Russia and Moldova to other former Soviet Republics and several European countries. Using local branching index, we found that this clade is expanding more rapidly than comparable clades of lineage 4.2. This evolutionarily successful strain’s presence beyond Moldova poses a threat to MDR-TB control, and enhanced surveillance is necessary to reduce the risk of MDR-TB epidemics in the region.

**Funding:** The authors report funding from the National Institutes of Health (R01AI180209, P01AI159402)

## Introduction

Multidrug resistant tuberculosis (MDR-TB) is an emerging global health challenge for TB elimination efforts. While drug resistance-conferring mutations can arise over the course of treatment, transmission of drug-resistant *Mycobacterium tuberculosis* (*Mtb*) strains sustains MDR-TB epidemics in high burden settings.^(1)^ Several recent studies have highlighted the role of MDR *Mtb* transmission in the Russian Federation and former Soviet Republics,^(2-4)^ where, in some settings, over 40% of new TB cases have a drug-resistant phenotype.^(5)^ Of the ten human-adapted *Mtb* lineages, the lineage 4 is among the most strongly associated with MDR phenotypes, and appears to be more likely to become the dominant multidrug resistant strains within local epidemics.^(6)^ Lineage 4 is genetically diverse and geographically widespread, supporting both globally represented “generalist” strains and geographically restricted “specialist” strains.^(7)^ The lineage 4.2/Ural strain has been described as an “intermediate” strain; strains belonging to this sublineage has been identified in several eastern European, central Asian, and east African countries,^(7)^ and may have the potential to spread widely.

Several recent studies have described a highly successful strain of MDR lineage 4.2/Ural *Mtb*. A study of lineage 4.2/Ural *Mtb* in eastern Europe identified an epidemic clone resistant to rifampin, isoniazid and kanamycin that had reached epidemic proportions in the Republic of Moldova within the last 25 years.^(8)^ A study in the Moldovan capital of Chisinau described a similar MDR lineage 4.2/Ural strain that emerged in the 1990s and underwent significant expansion over the same time period.^(9)^ Most recently, a country-wide *Mtb* phylogeographic analysis identified a large clade of the same Ural strain with evidence of high levels of transmission throughout Moldova.(^2, 10^) This strain was estimated to have an effective reproduction number twice that of drug-susceptible lineage 4.2 strains, and appeared to be expanding more rapidly than lineage 2 MDR strains in Moldova.^(11)^ Global data suggest that the emergence of multidrug resistant lineage 4 strains is a local phenomenon, with limited evidence of migration of resistant strains across borders.^(12)^ However, some Ural MDR-TB strains isolated in the Republic of Georgia^(4)^ appeared to be genetically similar to the rapidly spreading Ural MDR-TB strains in Moldova^(11)^, suggesting there may be more widespread dispersal of this lineage.

In this study, we constructed a global dataset of approximately 200,000 *Mtb* whole genome sequences available from public databases to assess the prevalence of lineage 4.2/Ural strains and identify strains genetically similar to the highly successful lineage circulating in the Republic of Moldova. Using genomic epidemiological analyses, we described the geographic spread, relative transmission fitness, and evolutionary history of this strain.

## Results

### Identification and global dispersion of lineage 4.2/Ural sequences

We analyzed 5909 *Mtb* whole genome sequences classified as lineage 4.2/Ural strains that were downloaded from the European Nucleotide Archive (ENA) (Supplementary Figure 1). The country of origin was available for 5440 sequences (92%) and the date of specimen collection was available for 4062 sequences (69%). We identified sequences from 61 countries across 6 continents (Figure 1). The countries with the largest share of lineage 4.2/Ural sequences were the Republic of Moldova (1546; 26%) and the Republic of Georgia (602; 10%). Notably, in several central-European countries we did not identify any lineage 4.2/Ural sequences in publicly available datasets. The completeness of date information varied: 1373 (23%) had complete dates, 2452 (41%) had partial dates, and 237 (4%) had a range of possible collection years. The oldest included sample was collected in 1994, and the newest included sample was collected in 2023.

**Figure 1:**
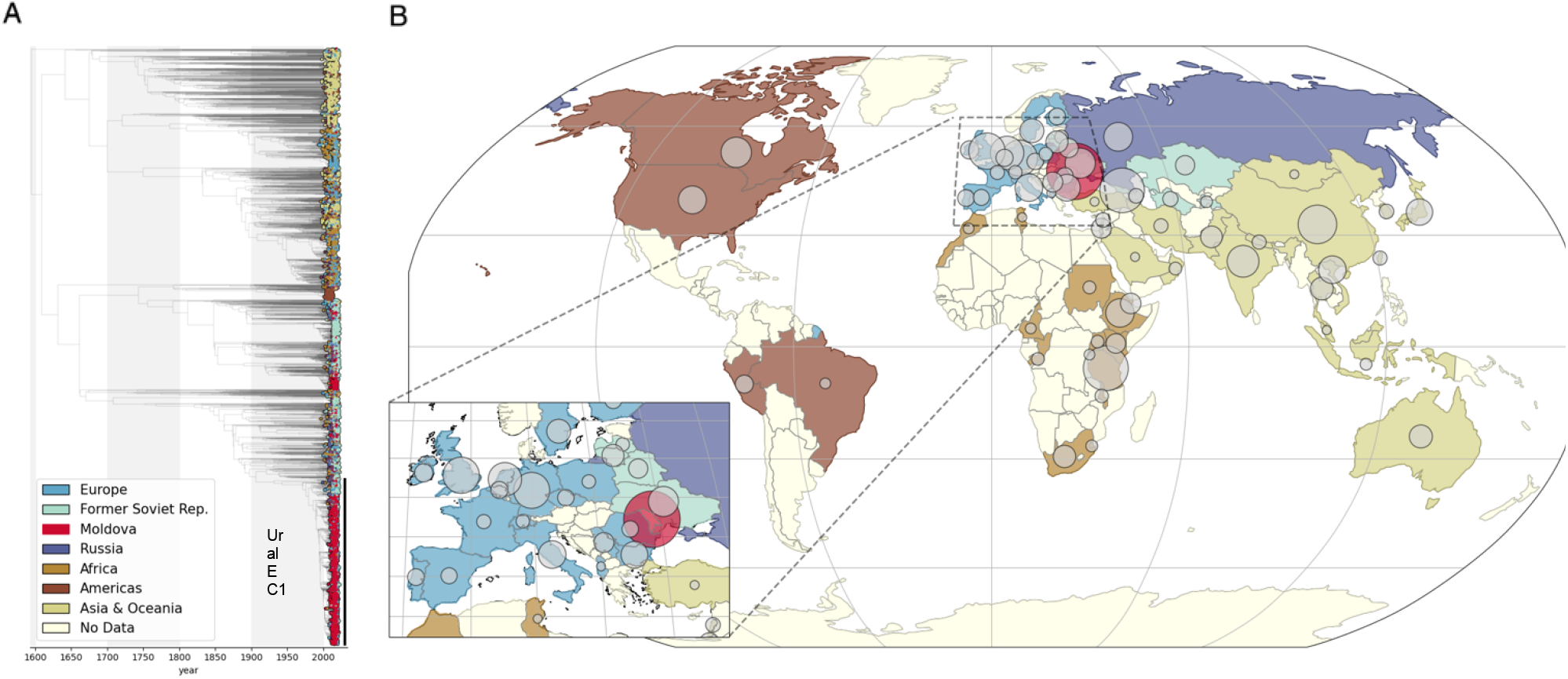
Global Dispersion of lineage 4.2 *Mtb* strains. (A) Time-calibrated phylogeny containing 5909 *Mtb* sequences included in the study; taxa are colored according to sequence region of origin. Ural EC1 is marked with a vertical black line. (B) Map of countries of origin for included sequences; circle is proportional to the number of sequences included from each country.

### Identification and dispersion of Ural EC1

We inferred a maximum-likelihood phylogeny (Supplementary Figure 2) and subsequently constructed a time-calibrated phylogeny by time-scaling branches using sampling dates. We identified a large clade (n = 1604) of MDR *Mtb* sequences that harbored clade-defining mutations (70 SNPs and 5 small insertions and deletions indels)^(11)^ and were genetically similar to the previously identified MDR *Mtb* strain in Moldova(^2, 8, 9, 11)^ (Figure 2). Strains with the same genetic background have previously been called Ural Clade C,^(8)^ multidrug resistant outbreak strain,^(9)^ Ural Clade 1,^(2)^ and Ural_A.^(11)^ Here, we refer to this clade as Ural epidemic clade 1 (EC1).

**Figure 2:**
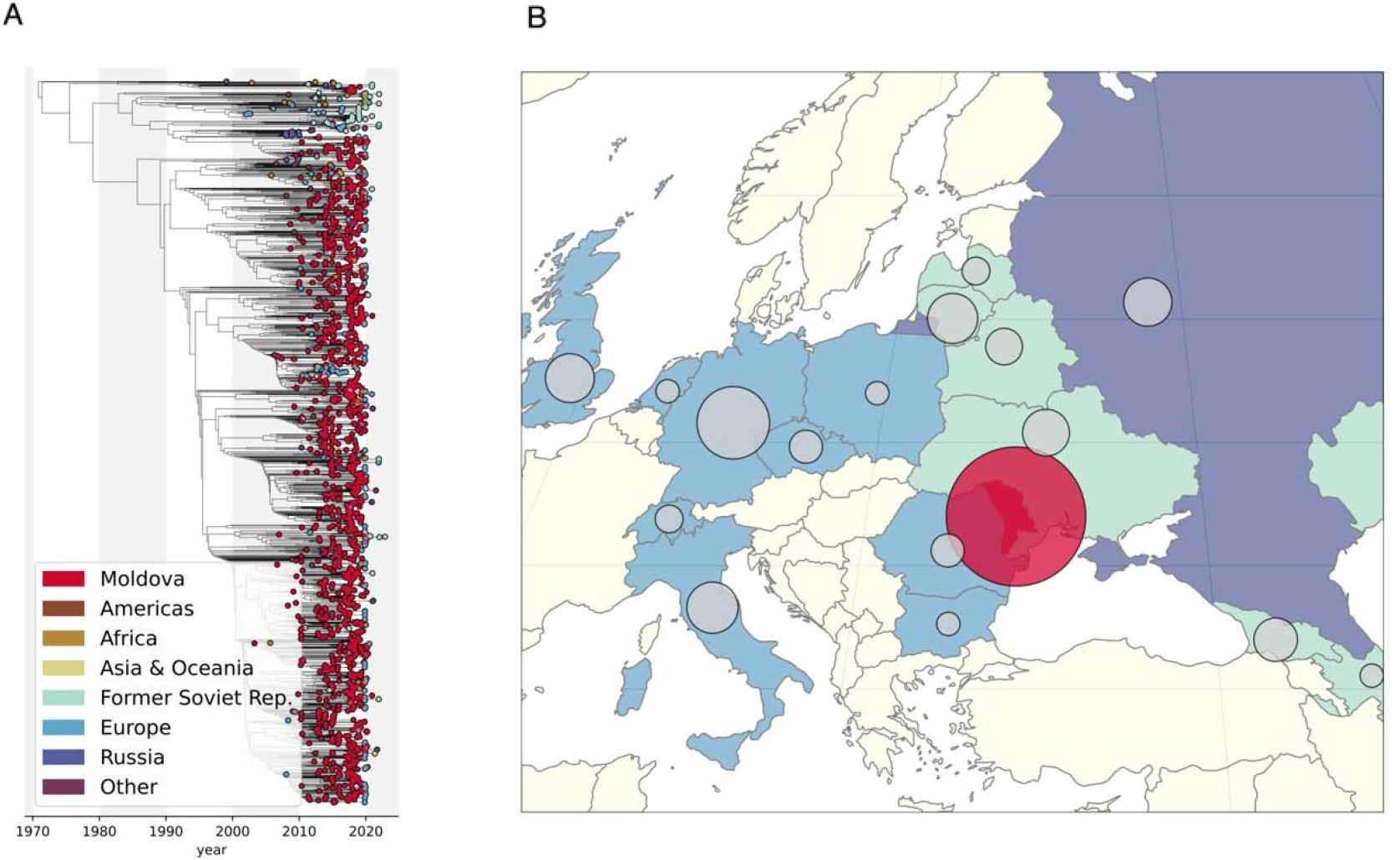
Ural EC1 Phylogeny and Geographic Dispersion. (A) Time-calibrated phylogeny containing 1604 *Mtb* sequences identified as part of Ural EC1; taxa are colored according to sequence region of origin. (B) Map of countries of origin for sequences in Ural EC1; circle is proportional to the number of sequences included from each country.

All Ural EC1 isolates had mutations associated with isoniazid (INH) resistance. Additionally, 1528 (95%) also carried mutations associated with rifampin (RIF) resistance and thus were MDR; this group included 399 pre-extensively drug resistant (XDR) isolates (MDR plus resistance to fluroquinolones [FLQ]) and 16 XDR isolates (pre-XDR plus resistance to one Group A drug, e.g., bedaquiline [BDQ] or linezolid [LZD]). The country of collection for sequences in the Ural EC1 clade was predominately Moldova (n = 1256; 78%), with sequences from Russia and other former Soviet republics comprising 5% of the dataset (n = 78) and 10% from other European countries (n = 155). Based on the time of most recent common ancestor (tMRCA) of the timed clade, we estimated that Ural EC1 emerged in 1971 (95% CI: 1965, 1976).

To characterize the movement of this strain across national borders, we inferred the country of internal nodes of the phylogeny using Sampling Aware Ancestral State Inference (SAASI), an ancestral state inference method that explicitly accounts for sampling differences (Figure 3). We infer that Ural EC1 emerged first in Russia (root state probability = 0.98) and that Russia was the source country for 128 migration events (45%). Migration events from Russia to Moldova accounted for 73% of outflow from Russia and 72% of inflow to Moldova. We inferred a relatively small number of migration events into other countries of the former Soviet Union; of the 31 events, Russia was the source country for 17 (55%) events and Moldova was the source country for 6 (19%). Moldova was the inferred country of origin in 96 total migration events; the majority (84 events, 88%) were migrations from Moldova into countries in Europe, primarily Germany (Supplementary Figure 3A).

**Figure 3:**
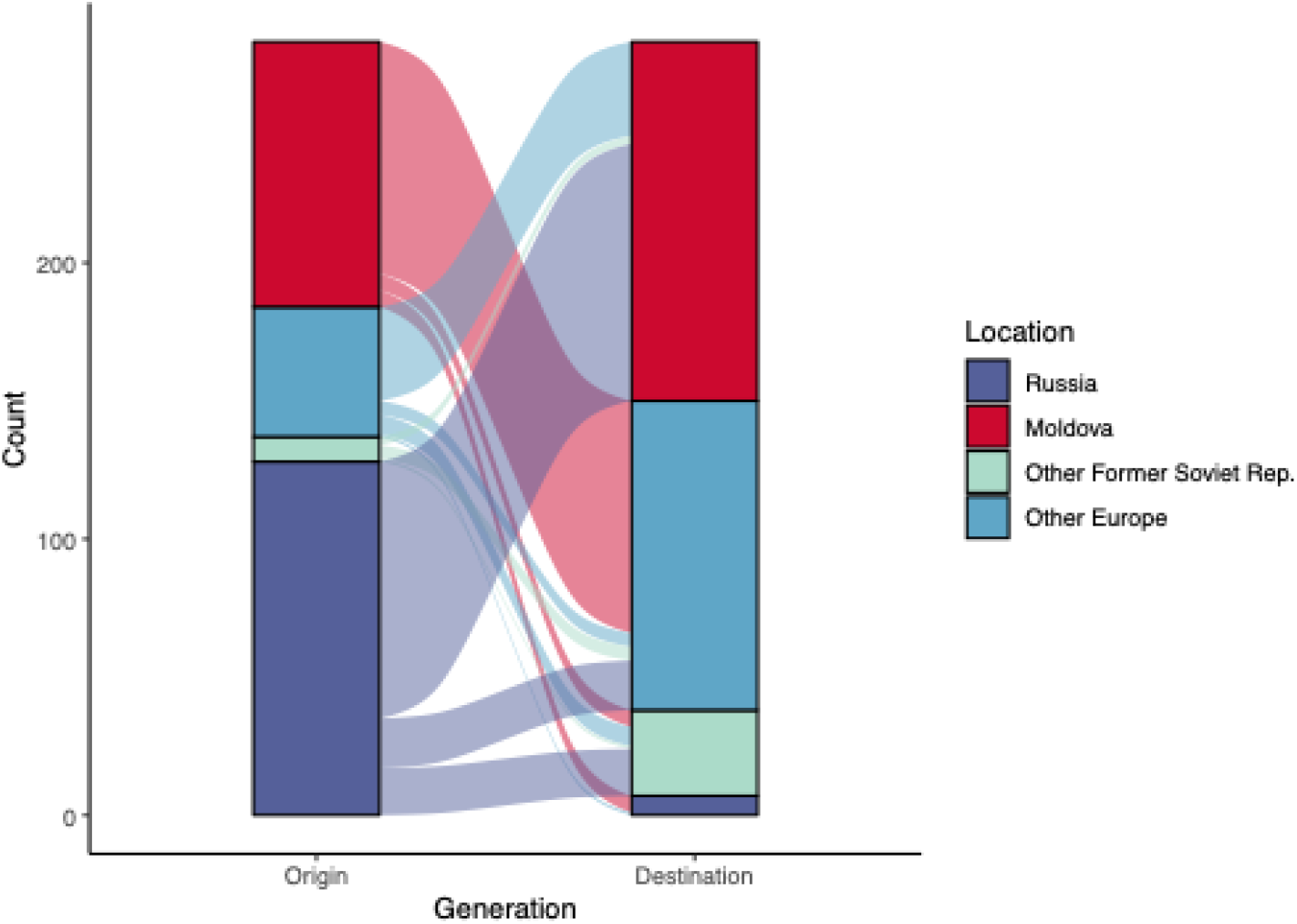
Migration of Ural EC1 strains. Alluvial plot showing the inferred origin and destination for each migration event. Sequences from countries with fewer than five isolates have been excluded. “Other Former Soviet Rep.” includes Belarus, Georgia, Lithuania, and Ukraine; “Other Europe” includes Germany, Italy, Portugal, and the United Kingdom. Migration events between countries within these groups have been included in the plot.

As a sensitivity analysis, we performed a conventional ancestral state reconstruction using the R package *ape* (which does not account for the sampling variability present in the data).^(13)^ Even when we ignored sampling variability, we found that Russia was still the most likely source of Ural EC1 (root state probability = 0.71). As expected, the method inferred a higher rate of migration events from Moldova (Supplementary Figure 3B).

### Evolution of drug resistance in the EC1 clade

We used ancestral state reconstruction to predict the emergence and distribution of 24 key drug resistance-conferring mutations to eight antimicrobials (RIF, INH, ethambutol [ETH], FLQ, kanamycin [KAN], streptomycin [STR], LZD), as well as RIF compensatory mutations (Figure 4). All Ural EC1 isolates carried the *kat*G Ser315Thr mutation that confers resistance to INH. Of those, 1442 (90%) also had an *inh*A .-777C>T mutation (also referred to as *fabG1* .-15C>T) in the upstream regulatory region of the *fabG1*-inhA operon, which confers low-level resistance to INH and was predicted to have emerged in the mid 1980s. A smaller number of other isolates (84; 5.2%) instead carried the *inh*A .-154G>A mutation in the same regulatory region, which emerged in the late 1980s.

**Figure 4:**
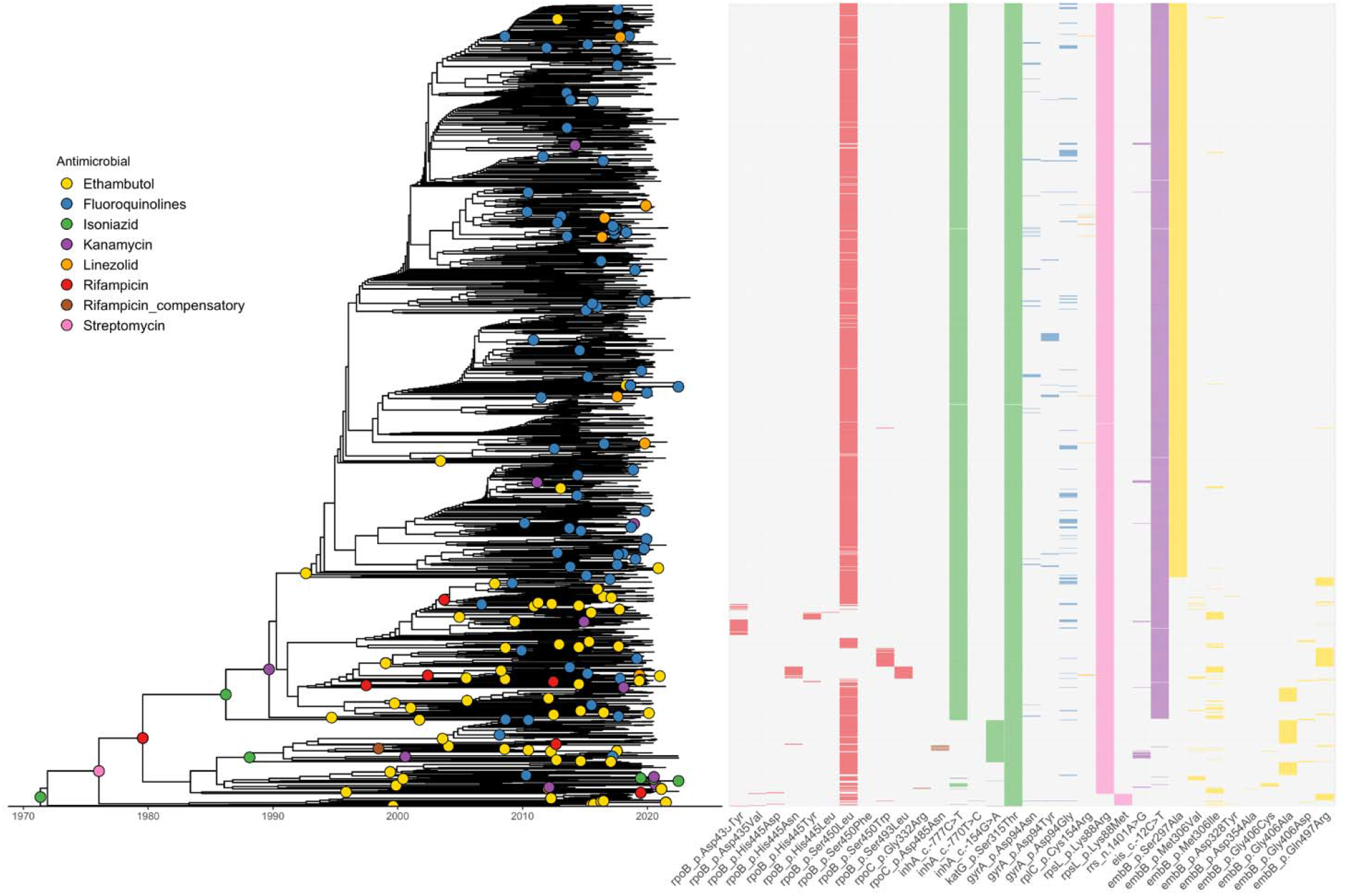
Ancestral state reconstruction of drug resistance-conferring mutations. Phylogenetic tree of Ural EC1 with the inferred drug resistance genotypes.

Most isolates (1406; 88%) harboured the *rpoB* Ser450Leu mutation conferring RIF resistance, which first emerged in Ural EC1 in the late 1970s. Of the remaining isolates, 123 contained a single other RIF resistance-associated mutation in *rpoB* and 21 strains had a double mutation in *rpoB* His445Asn and Ser493Leu that conferred RIF resistance. These mutations emerged several times from the late 1990s onwards. There were also 12 isolates that had mutations in *rpo*C that have previously been associated with compensatory mechanisms to RIF resistance (*rpoC* Gly332Arg^(14)^ and Asp485Asn^(15)^) and first emerged in the late 1990s in the clade.

We infer that resistance to STR first evolved in the clade in the mid 1970s as a result of the *rpsL* Lys88Arg mutation, which was carried by 1579 isolates (98%); 23 of the 25 remaining isolates later acquired the alternative *rpsL* Lys88Met mutation at the same loci. From the mid 2000s onwards, there were many occurrences of three FLQ resistance-conferring mutations evolving independently in the *gyrA* gene. Additionally, 1427 (89%) isolates carried the *eis* .-12C>T mutation—associated with resistance to KAN—that first emerged around 1990, with 14 of these also possessing the *rrs* .1401A>G KAN resistance-conferring mutation and 14 additional KAN resistant isolates carrying only this *rrs* mutation. Finally, we found that resistance to newer antimicrobials was uncommon in Ural EC1. Recent evolution of LZD resistance was identified in 11 isolates that carried the *rplC*Cys154Arg mutation, and four isolates contained a duplication in *mmpR5* that is associated with BDQ resistance.^(16)^

### Recent Expansion

We compared the distribution of the local branching index (LBI)^(17)^ of taxa in Ural EC1 to taxa in comparison clades. LBI is a measure of relative transmission fitness based on the topology of the phylogenetic tree. High fitness ancestors (internal nodes) will produce more rapid branching patterns in the phylogeny, and sampled isolates (taxa) of higher reproductive fitness can be identified as their recent descendants.

Four other lineage 4.2/Ural clades with at least 150 taxa and a tMRCA within 50 years of the emergence of Ural EC1 were used for comparisons: clade 1 (tMRCA = 1960 [1954, 1967], n = 212), clade 2 (tMRCA = 1949 [1945, 1953], n = 251), clade 3 (tMRCA = 1987 [1982, 1991], n = 185), and clade 4 (tMRCA = 1945 [1937, 1952], n = 152) (Supplementary Figure 4). We found that Ural EC1 had a higher median LBI than the taxa in other clades (Tukey test p value < 0.001 for all comparisons to other clades) (Figure 5a). We also found that, across countries, Ural EC1 had a higher LBI on average than taxa in comparison clades (Figure 5b).

**Figure 5:**
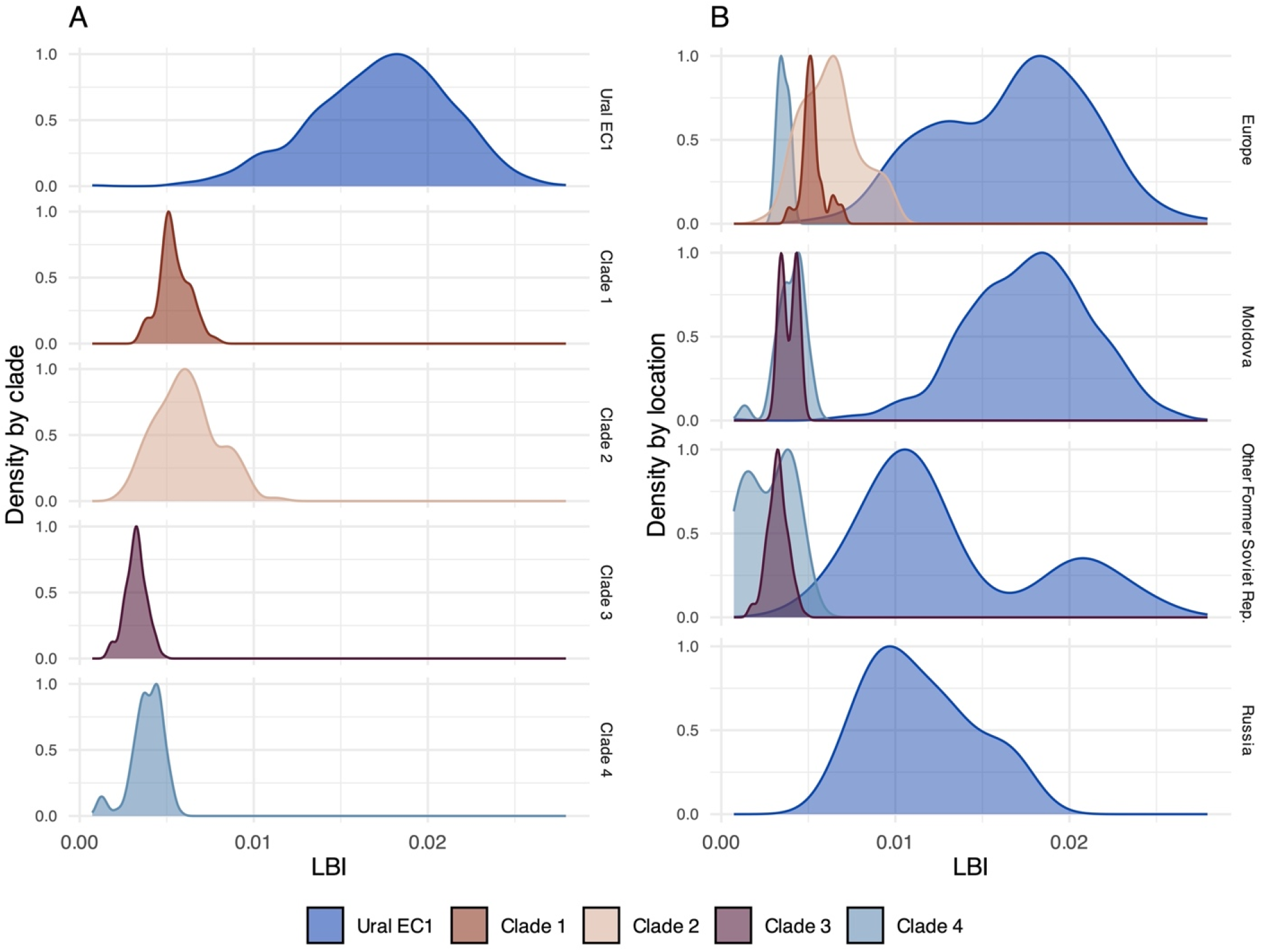
Local Branching Index. (A) Distribution of LBI for taxa in Ural EC1 and comparison clades. (B) Distribution of LBI for taxa in Ural EC1 by country or region of origin, colored by clade.

Within Ural EC1, the LBI for isolates from Russia did not differ significantly from those from former Soviet republics (excluding Moldova; Tukey test p value = 0.48). However, those from Moldova had a higher average LBI than isolates from other former Soviet republics (Tukey test p value < 0.001) and from Russia (Tukey test p value < 0.001). In the former Soviet Republics (excluding Moldova), we observed a bimodal distribution of LBI within Ural EC1. Most sequences in this group come from Georgia, Lithuania, and Ukraine; Ukraine had a higher LBI on average than Lithuania (Tukey test p value = 0.002).

## Discussion

We identified almost 6000 *Mtb* sequences belonging to the *Mycobacterium tuberculosis* complex lineage 4.2/Ural in publicly available sequence repositories. Among those sequences, 1604 belonged to a large clade of drug-resistant *Mtb* that contains MDR *Mtb* strains from Moldova with a high effective reproduction number.^(11)^ Using LBI, we estimated that this clade, Ural EC1, is growing more rapidly than other *Mtb* lineage 4.2/Ural clades emerging over similar time periods.

Our analysis suggested that Ural EC1 likely emerged in Russia around 1971 and subsequently spread throughout Europe and central Asia. This finding differs from an earlier analysis that suggested similar strains emerged in Moldova and spread to neighboring countries, including Georgia and Russia.^(9)^ In that study, the authors included a limited number of sequences from outside of Moldova and identified Moldova as the country of origin with only moderate certainty (posterior probability = 0.66). Our analysis included a larger sample of isolates from a broader set of countries, and we identified Russia as the country of origin with a high degree of certainty (probability = 0.98). Both studies conclude that there has been subsequent spread of this strain from Moldova into neighboring countries. However, our analysis suggests that migration events out of Russia also played an important role in the international spread of Ural EC1.

We found evidence that most MDR isolates in Ural EC1 carried the same *rpoB* mutations and all had a fixed *katG* mutation that confer resistance to rifampin and isoniazid, respectively. Ancestral state reconstruction suggested these mutations first evolved in the 1970s and were conserved in subsequent generations. Conversely, we found that the evolution of fluroquinolone resistance was likely driven by recent, independent acquisitions of mutations in the *gyrA* gene. Rifampin and isoniazid are first-line drugs that have been used to treat TB for decades; mutations which confer resistance to these first-line drugs have had many years of selective pressure pushing them towards fixation. Fluroquinolones are newer, second-line drugs used to treat individuals with MDR-TB, and there have been fewer opportunities for positive selection to favor mutations which confer resistance to these antimicrobials.

We attempted to include every publicly available lineage 4.2 *Mtb* sequence from ENA. This allowed us to describe the global distribution of lineage 4.2 strains and identify migration events in greater detail than in previous analyses.^(12)^ In some settings, isolates were collected as part of a city- or country-wide prospective whole genome sequencing study,^(15, 18)^ though in other settings sequences were part of dedicated studies on MDR-TB.^(19, 20)^ Because of the variable sampling strategies, it is challenging to fit phylodynamic models to these data. In the case of the country-level ancestral state reconstruction, we were able to overcome this challenge by using an approach that adjusts for heterogenous sampling. However, this still presents a limitation when determining the extent to which MDR lineage 4.2 isolates are present outside of Moldova, where there have been several large sequencing studies.^(2, 8, 9)^

We were also limited by the availability of strain metadata; for example, data was not available on the country of origin for 8% of isolates or the date of isolation for 31% of isolates. Finally, several European countries had no available lineage 4.2 *Mtb* sequences. In many of these settings, we are aware of the existence of *Mtb* whole genome sequence data that have not been made publicly available (e.g. EuSeqMyTB).

We present new evidence to suggest that a rapidly expanding strain of MDR *Mtb*, which was previously believed to be restricted to the Republic of Moldova, has spread to other European and central Asian countries. The broad dispersal of Ural EC1 is an urgent threat to TB control in the region. While rapid molecular tests can quickly identify drug resistant disease, they cannot identify specific bacterial lineages or strains. Routine whole genome sequencing is therefore essential to support surveillance of Ural EC1 and other strains of concern.

### Online Methods

#### Global collection of Mtb Ural lineage 4.2 whole genome sequence data

We queried ENA on 18 February 2024 for all *M. tuberculosis* genomes (n = 196,547 accessions). ENA is synchronized with GenBank, making it the most complete public source of whole genome sequencing data for *Mycobacterium tuberculosis* complex (MTBC). We excluded laboratory and reference strains, other Mycobacteria species in the MTBC, and samples isolated from non-human hosts. We compared the remaining sample accession numbers (n = 177,856) in ENA to the TB-Profiler (TBP) database,^(21)^ which contains lineage assignments and drug resistance predictions. All sequences without an established lineage or sub-lineage assignment in the TBP database (n = 41,233) were subsequently profiled. We identified 7165 unique sample accessions that were profiled as *Mtb* lineage 4.2, which comprised 7563 whole genome sequencing data files (including samples with duplicate sequencing data or that were re-sequenced).

The 7563 sequencing data files were downloaded from ENA and aligned to the H37Rv reference strain (NC_000962.3) using BWA-MEM^(22)^ for both paired and single end read data. Binary alignment (BAM) files were indexed and sorted with SAMtools.^(23)^ Alignments with less than 80% mapping to the H37Rv reference strain and an average read depth below 50x were removed, along with any sample with evidence of mixed infection detected using MixInfect2.^(24)^ In cases where samples had multiple run accessions (duplicate or re-sequenced isolates), alignments with the highest mapping and average read depth were retained for a final dataset of n = 5909 *Mtb* lineage 4.2 sequences (one clinical sequence per sample specimen) (Supplementary Data 1).

Variant calling was conducted using GATK^(25)^ ‘HaplotypeCaller’ and ‘GenetypeGVCFs’; low-confidence variants (Q < 20, read depth < 5) and sites with an ambiguous or missing call in more than 10% of isolates were removed. The consensus nucleotide (≥80% of mapped reads) was assigned at loci with mixed calls, otherwise the nucleotide ‘N’ was assigned. Finally, variants in repetitive regions, in PE/PPE genes, and at known resistance-conferring loci, were removed. A multi-sequence alignment of variant SNPs was constructed for subsequent analyses.

We cross-referenced sample country and collection date between ENA and TBP. In cases of conflicting country metadata, we preferentially used the value from ENA. For conflicting dates, we used the most complete date available; when both dates were complete, we used the date recorded in ENA. For isolates belonging to large projects (10 or more sequences included in this study) with missing collection country or date, we queried PubMed for publications associated with the BioProject ID. If metadata were not available as a supplement to these studies, we requested these data from corresponding authors.

#### Phylogenetic reconstruction

We performed maximum-likelihood phylogenetic reconstruction using *IQ-TREE 2*^(26)^ from a multi-sequence alignment of concatenated SNPs. The optimal substitution model (TVM+F+G4) was determined using the model test (‘-m’) option, and branch support was calculated using 1000 bootstrap replicates. We then performed Bayesian inference of a time-calibrated phylogenetic tree using the R package *BactDating*,^(27)^ fitting the model using the maximum-likelihood phylogeny after scaling the branch lengths to SNPs/genome/year, and calibrating the tree using sampling dates. When sampling dates were uncertain or unavailable, we used uniform priors with bounds indicating the earliest and latest possible sampling date. We fit the model using a fixed mean clock rate of 0.5 SNPs/genome/year (approx. 1.145×10^-7 SNPs/site/year) and a strict gamma clock model. We ran the model for 5 × 10^5^ MCMC samples, thinning the posterior by a factor of 500, resulting in 1,000 posterior samples.

We characterized a monophyletic clade within the time-calibrated phylogeny that included all isolates previously identified as part of a rapidly expanding MDR-TB strain in Moldova that harbored clade-defining mutations; 70 SNPs and 5 small insertions and deletions (indels).^(11)^ Finally, we used the R package *treestructure*^(28)^ to identify comparable lineage 4.2 clades, each with 150 or more taxa and a tMRCA within 50 years of Ural EC1’s most recent common ancestor.

#### Phylodynamic and genomic analyses

We performed ancestral state reconstruction using SAASI, an ancestral state inference method that explicitly accounts for sampling differences and is computationally feasible on large trees. We used SAASI to infer the ancestral states of EC1 strains from Belarus (n = 6), Georgia (n = 12), Germany (n = 92), Italy (n = 23), Lithuania (n = 21), Moldova (n = 1256), Portugal (n = 8), Russia (n = 18), Ukraine (n = 16), and the United Kingdom (n = 19).

SAASI requires estimates of transition, speciation, extinction, and sampling rates. We estimated the transition rates between different countries using the ape package in R.^(13)^ We specified a three-parameter model: (1) a transition rate from Moldova to other countries (estimate: 0.001), (2) a transition rate from other countries to Moldova (estimate: 0.009), (3) a transition rate from any pair of the non-Moldova countries (estimate: 0.004). Next, we estimated the speciation (estimate: 0.174) and extinction (estimate: 0.001) rates using a maximum likelihood approach,^(29)^ assuming that the sampling rate is known. Finally, we estimated sampling rates by first estimating sequencing coverage by country and then scaling that to the average sequencing coverage in the clade and the inferred speciation rate. We use the following equations:

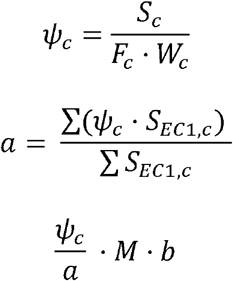

where *S*_*c*_ is the observed number of lineage 4.2 sequences, *W*_*c*_ is the estimated number of TB cases using WHO notification data, *F*_*c*_ is the estimate fraction of cases belonging to Lineage 4.2. We estimate this fraction using data from 2015-2019, the same period in which 50% of the sequences in the clade were collected. We assumed 10% sequencing coverage when data on TB incidence were unavailable, and we assumed no country had a sequencing coverage > 50%. We normalize ψ_*c*_ by a, the weighted average sequencing coverage in the Ural EC1 clade. Since this results in a fraction, not a rate per unit time, we multiply this value by the inferred speciation rate *M*, scaled by a factor b (chosen such that no sampling rate exceeds the inferred speciation rate; b = 0.8).

The emergence of key drug resistance mutations in Ural EC1 was inferred using a maximum likelihood marginal reconstruction of ancestral sequences at nodes in the timed phylogeny, implemented in the R package *Phangorn*.^(30)^ We included ambiguous sites and missing calls to reflect prior probabilities of all character states. Mutations conferring resistance to RIF, INH, ETH, FLQ, KAN, STR, and LZD, along with rifampin resistance compensatory mutations, were determined using the WHO catalogue.^(31)^

Finally, we calculated LBI^(17)^ at every node in the maximum likelihood phylogeny of lineage 4.2 (n = 11817, taxa = 5909) using a neighborhood size of 2.18 × 10^−4^ (0.0625 times the average pairwise patristic distance [3.5 × 10^−3^ substitutions/site]). We report the distribution of LBI for each terminal node. We compare the distribution of LBI across groups using Tukey’s test for multiple comparisons.

This manuscript follows the STROME-ID guidelines.^(32)^

## Notes

### Competing Interest Statement

The authors have declared no competing interest.

