## Supplementary Figures for "The global phylogeography of rapidly expanding multidrug resistant Ural lineage 4.2 *Mycobacterium tuberculosis*"

**Supplemental Materials**

Supplemental Data 1: Run accession, country of origin, and estimated sampling date for each sequence in Ural EC1.

Supplemental Figure 1: Inclusion criteria for sequences used in this study

Supplemental Figure 2: Maximum-likelihood of phylogeny with all 5909 isolates included in the study

Supplemental Figure 3: Ancestral state reconstruction

Supplemental Figure 4: Time-calibrated phylogeny with comparison clades

Supplemental Figure 1: Inclusion criteria for sequences used in this study

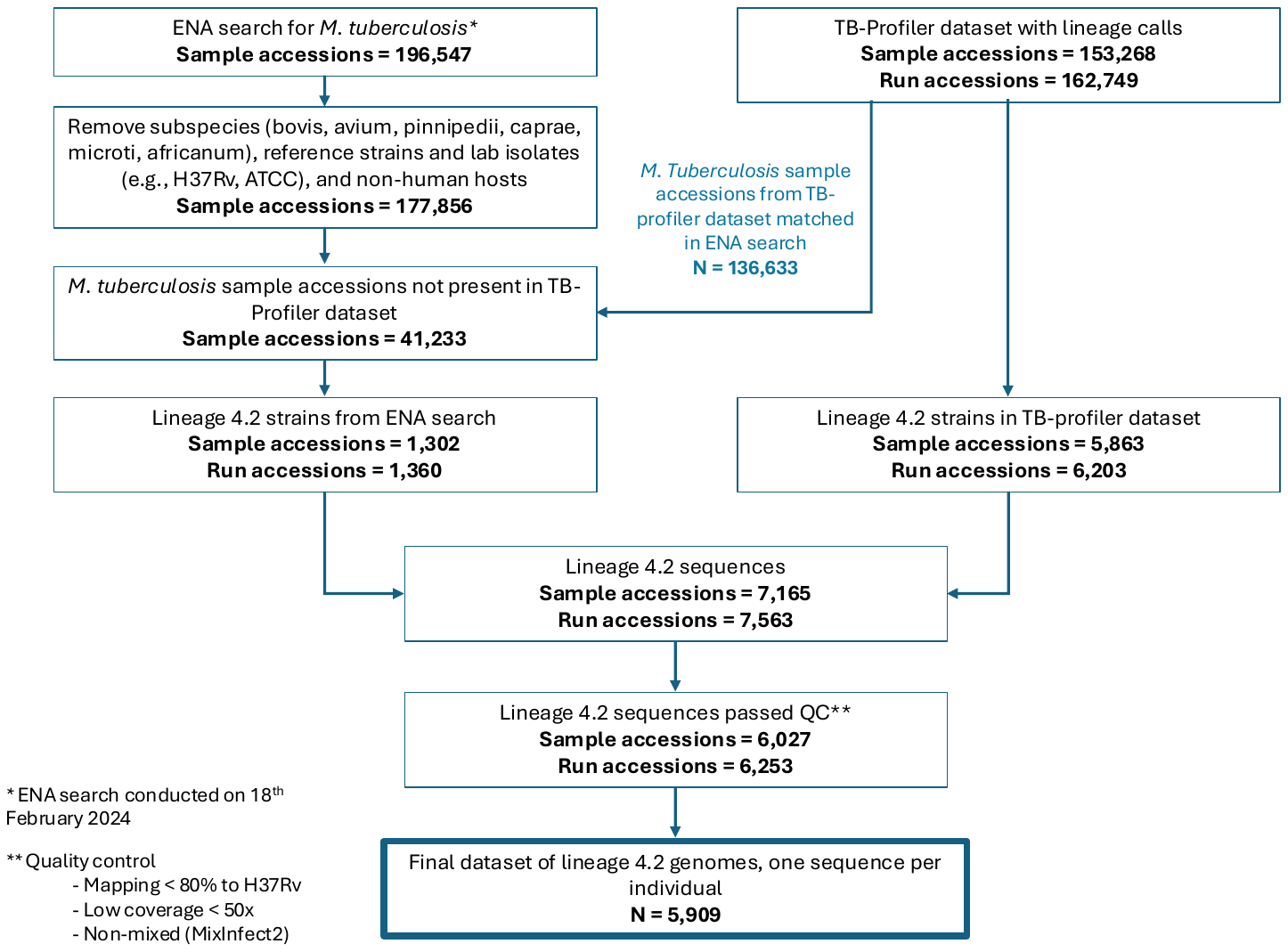

Supplemental Figure 2: Maximum-likelihood of phylogeny with all 5909 isolates included in the study. Strains belonging to EC1 are colored in blue; all other strains are colored in grey.

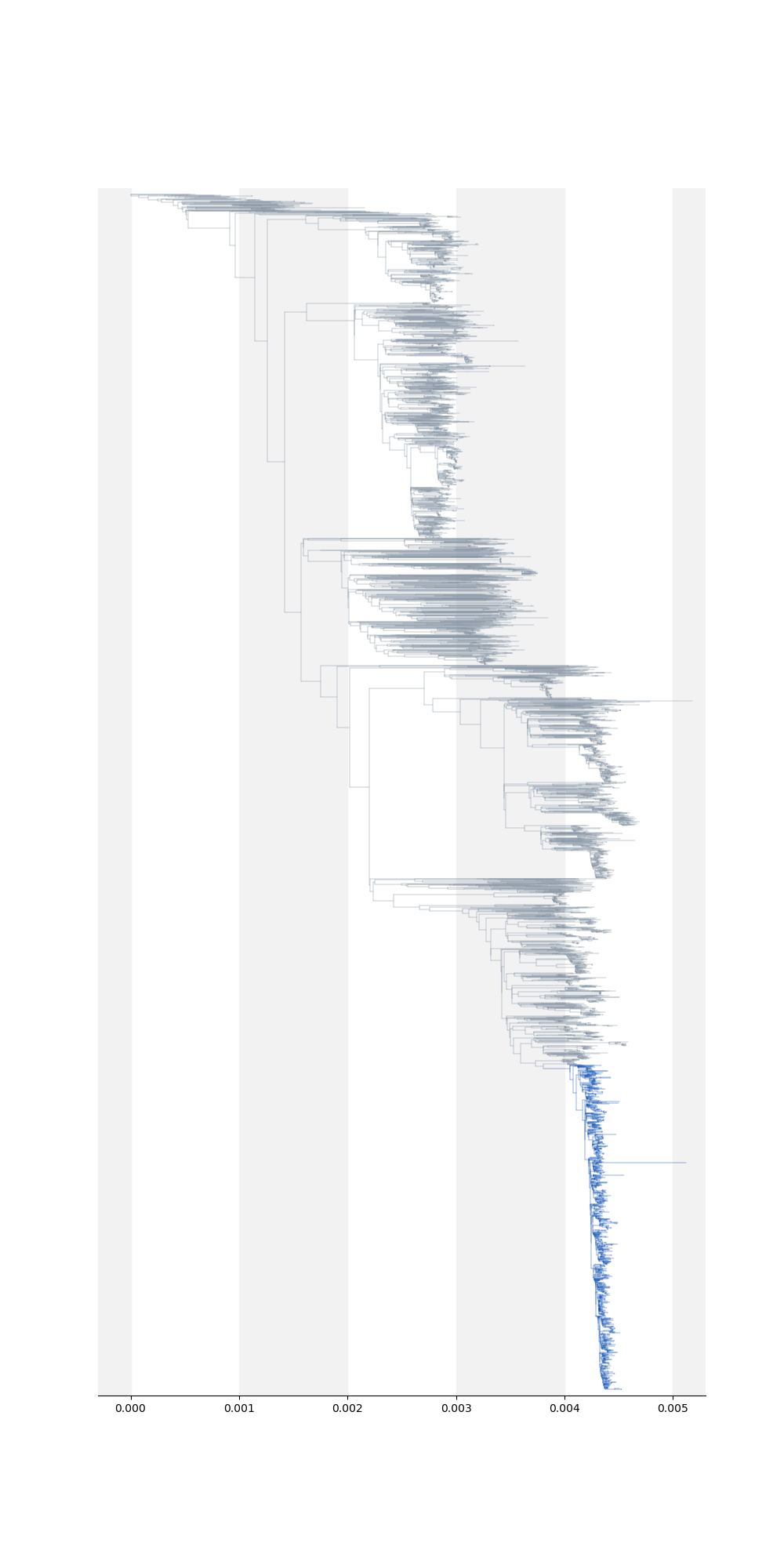

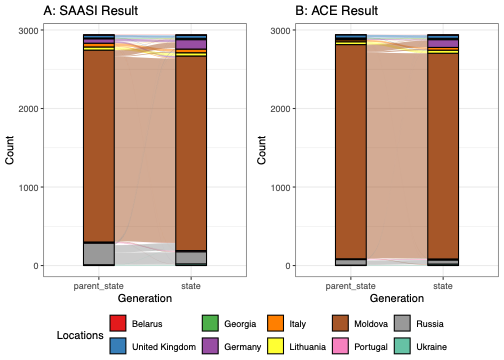
Supplemental Figure 3: Ancestral state reconstruction. (A) Result from SAASI model, including the inferred state of every node pair. (B) Result from ACE model, including the inferred state of every node pair.

Supplemental Figure 4: Time-calibrated phylogeny with comparison clades

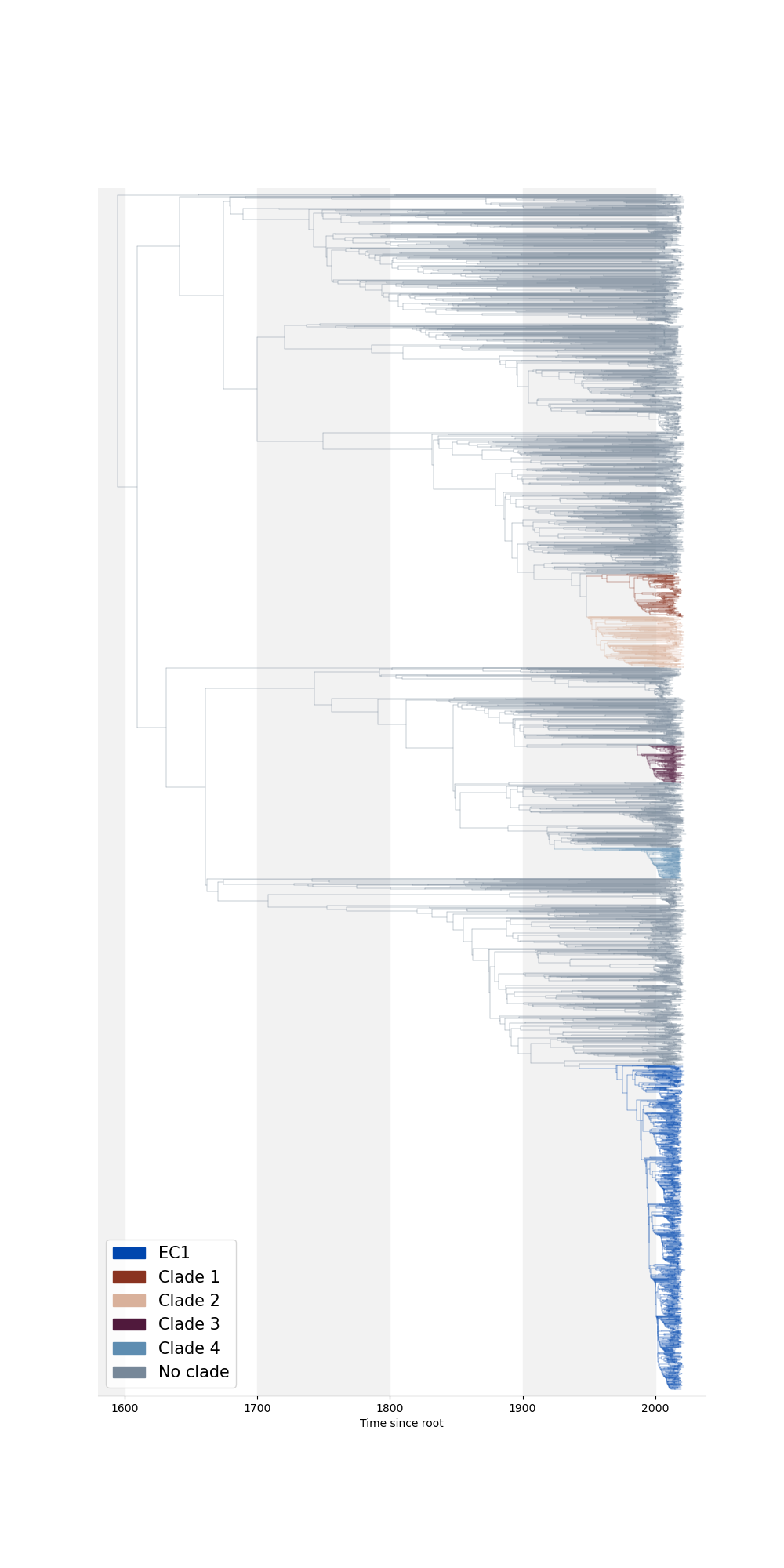
